# Effector loading onto VgrG spike proteins is critical for the assembly of the type VI secretion system in *Agrobacterium tumefaciens*

**DOI:** 10.1101/556712

**Authors:** Yun-Wei Lien, Chih-Feng Wu, Devanand Bondage, Jer-Sheng Lin, Yu-Ling Shih, Jeff H. Chang, Erh-Min Lai

**Author notes:** The first two-authors contributed equally to this work (Yun-Wei Lien, Chih-Feng Wu). Correspondence: Dr. Erh-Min Lai, Institute of Plant and Microbial Biology, Academia Sinica, 128, Sec. 2, Academia Road, Nankang, Taipei, Taiwan 11529.

## Abstract

The type VI secretion system (T6SS) is used by many bacteria to engage in social behaviors with others and can directly or indirectly affect the health of plants and animals. Because activities associated with T6SS are often costly, the assembly and activation of the T6SS must be highly regulated. However, our knowledge regarding how T6SS assembly and contraction are regulated remains limited. Here we show that the loading of effectors onto their cognate carriers is critical for the assembly of a functional T6SS in *Agrobacterium tumefaciens. A. tumefaciens* strain C58 encodes one T6SS and two Tde DNase toxin effectors used as major weapons for interbacterial competition. We found that loading of Tde effectors onto their cognate carrier, the VgrG spike, is required for active T6SS secretion. Our data also suggest the assembly of the TssBC contractile sheath occurs only after Tde effectors are loaded onto the VgrG spike. The requirement of effector loading for efficient T6SS secretion was also validated in other *A. tumefaciens* strains. Such a mechanism may be used by bacteria as a strategy for efficacious T6SS firing. Given the prevalence of T6SS-encoding loci in host-associated bacteria, these findings inform on mechanisms that influence the composition of microbial communities and the services provided to hosts.

## Introduction

The Type VI secretion system (T6SS) is a versatile secretion system that has been implicated in virulence, antagonism, nutrient acquisition, and horizontal gene transfer ^1–3^. Contact-dependent interbacterial competition appears to be the major function of T6SS, a function that can influence the composition of microbial communities ^4,5^ A T6SS machine resembles a contractile phage tail-like structure ^6–8^. Effectors are loaded onto the T6SS either via non-covalent interactions or as carboxy-terminal extensions to either of the three core structural components (Hcp, VgrG, or PAAR) ^2,9^ Upon contraction, the puncturing device carrying effectors is fired and propelled, carrying or allowing effectors, across the cell membrane into the extracellular milieu or into targeted prokaryotic or eukaryotic cells. In general, effectors are considered as accessary components of the secretion apparatus. This is based on the repeated observations that Hcp and/or VgrG, key markers for secretion, can be detected in the extracellular milieu of mutants lacking effector genes ^10–14^

The plant pathogenic bacterium *Agrobacterium tumefaciens* strain C58 encodes one T6SS main cluster consisting of the *imp* and *hcp* operons and *vgrG2* operon distal to the main cluster ^15^. Three T6SS toxin effectors were identified, in which secretion of Tde1 and Tde2 DNases is governed specifically by VgrG1 and VgrG2, respectively, and secretion of Tae amidase is likely mediated by Hcp ^16,17^ Tde effectors are major weapons deployed by *A. tumefaciens* for interbacterial competition *in planta*^10^. Each of the effector genes is genetically linked to a cognate immunity gene, which form three toxin-immunity pairs (i. e. *tae-tai* and *tde1-tdi1* located in the main gene cluster and *tde2-tdi2* encoded downstream of *vgrG2* distal to the main cluster) (Fig. 1a). Self-intoxication is prevented in the toxin-producing cells by the cognate immunity proteins.

**Fig. 1.**
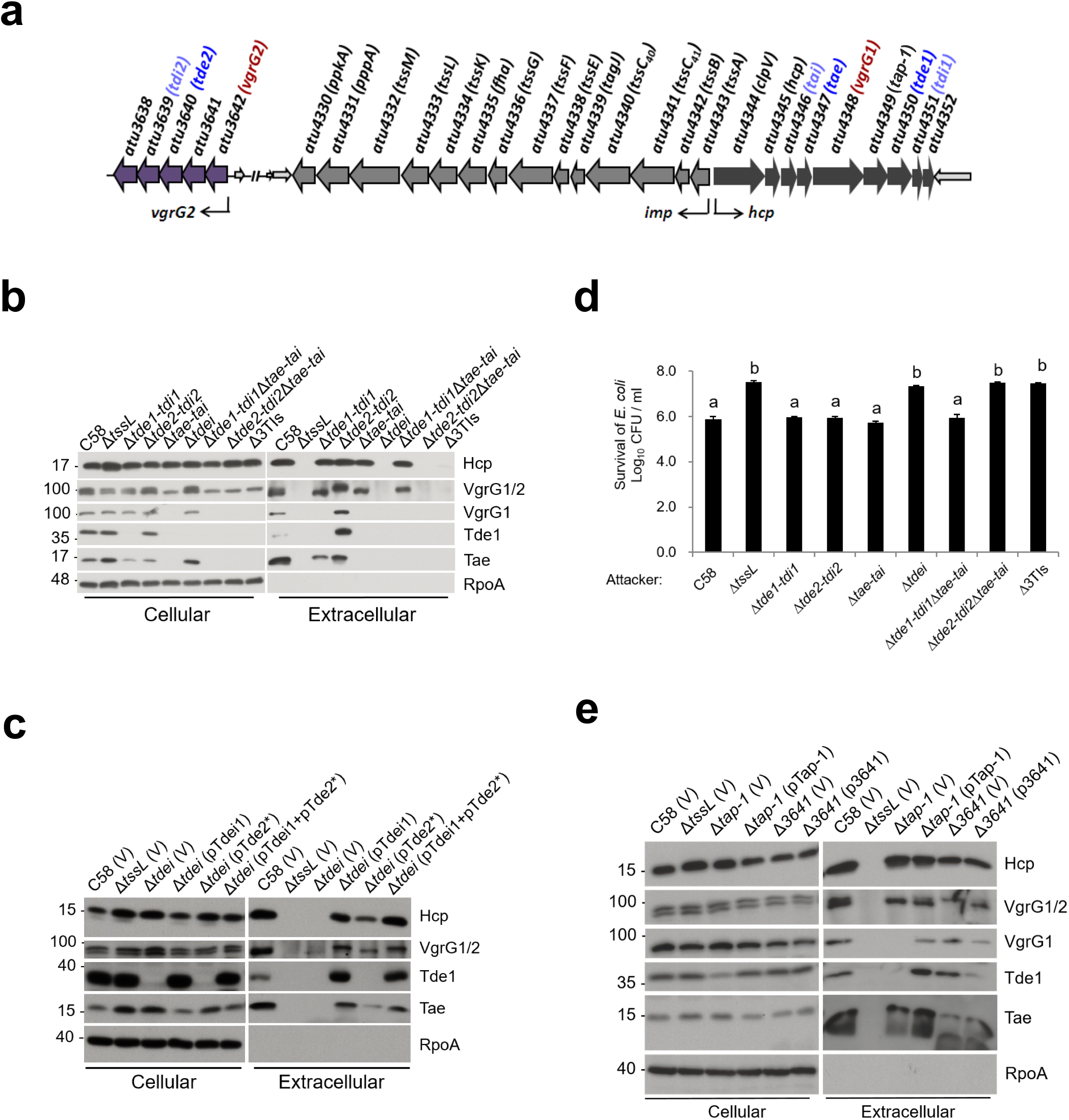
Tde effector loading onto VgrG is required for secretion of the cognate VgrG. (a) The structure of the *imp* and *hcp* operons and *vgrG2* operon in *A. tumefaciens* C58 ^10^. The arrows represent coding sequencing, with arrowheads depicting the direction of expression. The bigger arrows represent genes that code for components of the T6SS. The *vgrG2* operon is located distal to *imp* and *hcp*. The *vgrG* and toxin-immunity gene pairs are highlighted in colors. (b) T6SS secretion assay of *A. tumefaciens* strains: wild type C58, various mutants lacking one, two, or three toxin-immunity gene pairs, and a mutant lacking *tssL*. (c) T6SS secretion assay of various *A. tumefaciens* strains: wild type C58, Δ*tssL*, the *tde* double deletion mutant *(Δtdei)* containing pRL662 and pTrc200 empty vector (V) only or expression of pTdei1 *(tde1-tdi1* expressed from pTrc200), pTde2* (catalytic site mutated *tde2* expressed on pRL662), or pTdei1+ pTde2*. (d) *A. tumefaciens* antibacterial activity assay against *E. coli*. The strains of *A. tumefaciens* were co-cultured at a ratio of 30:1 with *E. coli* DH10B (+ pRL662) on LB agar. The survival of target *E. coli* cells was quantified by counting CFUs on gentamicin-containing LB agar plates. Data represent mean ± standard error (SE) of 3 biological replicates. Statistics were calculated and significant differences (P <0.01) are indicated with different letters. (e) T6SS secretion assay of *A. tumefaciens* strains: wild type C58, Δ*tssL, Δtap-1*, and Δ*atu3641* harboring a pTrc200 vector (V) or derivatives pTap-1 (*tap-1* expressed from pTrc200), or p3641 (*atu3641* expressed from pTrc200). Cellular and extracellular fractions were collected from *A. tumefaciens* strains grown in liquid 523 medium. Western blots were probed with indicated antibodies; the α-VgrG antibody detects VgrG1 (upper band) and VgrG2 (lower band) while α-VgrG1 detects only VgrG1 ^16^. RpoA is RNA polymerase, which is localized to the cytosol of *A. tumefaciens*. Molecular weight markers (in kDa) are indicated on the left.

Here we show that the loading of cargo effectors onto their cognate VgrG spike proteins is required for efficient T6SS-dependent secretion by *A. tumefaciens*. We demonstrate that in the absence of effector genes, the levels of secretion of Hcp and VgrG and formation of the TssBC sheath are significantly reduced in all examined strains of *A. tumefaciens*. Our study reveals a strategy that is deployed by bacteria to ensure that effectors are loaded onto T6SS prior to completing its assembly and firing outwards.

## Results and Discussion

Despite evidence suggesting that effectors are not components of the secretion apparatus ^10–14^, our previous study showed that variants of VgrG1 lacking the Tde1-binding domain is able to secrete but at slightly lower levels of Hcp and Tae effector ^16^. This led us to hypothesize that effector-loaded VgrG is more efficiently recruited for T6SS assembly and/or secretion. To test this hypothesis, we first examined whether the secretion of Hcp and VgrG proteins is affected by the presence or absence of effector genes. We found that the secretion of Hcp and VgrG proteins is affected by the presence or absence of effector genes in *A. tumefaciens*. In wild type C58, but not Δ*tssL*, a secretion-deficient mutant, Hcp, VgrG1/2, Tde1, and Tae were detected in the medium, referred as extracellular fraction, confirming their T6SS-dependent secretion (Fig. 1b). However, VgrG1 and VgrG2 proteins were not detected in the extracellular fraction of mutant strains in which their cognate effector gene was lacking. VgrG1 was not detectable in the extracellular fraction of any mutant minimally lacking the *tde1-tdi1* toxin-immunity gene pair (i. e. Δ*tde1-tdi1*, Δ*tdei* lacking both *tde1-tdi1* and *tde2-tdi2*, and Δ3TIs lacking all three toxin-immunity pairs). Similarly, VgrG2 is inferred to behave in a similar manner in mutants lacking *tde2-tdi2* toxin-immunity gene pair. Complementing the Δ*tdei* mutant with *tde1* restored its ability to secrete VgrG1 while the mutant expressing *tde2\** (encoding the secretable Tde2 variant with catalytic site mutation) failed to secrete VgrG1. In contrast, the mutant carrying *tde2\**, but not *tde1*, restored the secretion of VgrG2 (Fig. 1c). The levels of Hcp were similar in the extracellular fractions of the wild type strain as well as that from each of the mutants lacking a single toxin-immunity gene pair. During the course of this study, we noticed that in Δ*tae-tai*, downstream encoded proteins (VgrG1, Tap-1, and Tde1 proteins) were not detected in either of the cellular or extracellular fractions while *imp* operon-encoded TssB and ClpV encoded upstream of *tae-tai* were detected in all analysed strains (Fig. 1b and S1). This suggests that Δ*tae-tai* has a polar effect and explains why only VgrG2 but not VgrG1 were detected in Δ*tae-tai* and Δ*tde2-tdi2*Δ*tae-tai* mutants. Interbacterial competition assay showed that *A. tumefaciens* C58 can only kill *E. coli* when at least one Tde effector is delivered (Fig. 1d). The reliance on Tde DNases but not Tae amidase as the primary effectors against *E. coli* is consistent with previous finding that Tde but not Tae influence the *in planta* interbacterial competition activity of *A. tumefaciens*^10^.

Strikingly, the levels of secreted Hcp were hardly detectable in Δ*tdei*, Δ*tde1-tdi1*Δ*tae-tai*, and Δ3TIs. This result was surprising because in a previous study, the levels of secreted Hcp in the Δ3TIs mutant were similar to those of wild type C58 ^10^. A key difference between this previous study and the one presented here is that, in the previous study, the secretion assay was conducted in an acidic minimal medium (I-medium, pH 5.5), whereas here we used a rich medium for unambiguous detection of VgrG secretion. Therefore, we carried out secretion assay for strains Δ*tdei* and Δ3TIs in I-medium (pH 5.5). As reported previously, extracellular Hcp levels in Δ3TIs are similar to those from wild type C58. However, the levels of Hcp were low in Δ*tdei* (Fig. S2). Expression of Tae alone or the Tae-Tai pair in Δ3TIs reduced Hcp secretion levels while Tai expression in Δ3TIs did not impact levels of extracellular Hcp in Δ3TIs. These results suggested a role of Tae in regulating Hcp secretion levels in different growth conditions. Nevertheless, Hcp secretion levels are significantly reduced in Δ*tdei* grown in either growth condition. Therefore, we used the Δ*tdei* mutant to further investigate the mechanisms of effector-dependent secretion, because it consistently gave a secretion-less phenotype across different media/conditions.

Tde1 and Tde2 require specific adaptor/chaperone proteins to be loaded onto their cognate VgrG spike ^16^. Thus, we next examined whether secretion of the two VgrG spike proteins requires the cognate adaptor/chaperones. Secretion was assayed in Δ*tap-1* and Δ*atu3641*, mutants deleted of genes encoding the adaptor/chaperones for Tde1 and Tde2, respectively (Fig. 1e; (ref. 16)). Hcp, VgrG variants, Tde1 and Tae accumulated to detectable levels in the cellular fractions of all strains. VgrG1 and Tde1 were only detected in the extracellular fraction of cells that encoded Tap-1. Likewise, VgrG2 was detected in the extracellular fraction of cells that encoded Atu3641. In contrast, Tae remains detected in the extracellular fraction of cells in the absence of either of the adaptor/chaperone-encoding genes. In our previous study, when VgrG1 variants were expressed at higher than endogenous levels of wild type in a *vgrG1vgrG2* double deletion mutant (ΔG1ΔG2), variants of VgrG1 lacking the Tde1-binding domain are still able to mediate secretion of Hcp and Tae effector albeit at slightly lower levels ^16^. We then predicted that overexpression of VgrG in the absence of a cognate effector may be sufficient to initiate the assembly of the T6SS. The polymutant strain, Δ*tde1-tdi1*Δ*G1*Δ*G2op*, was generated lacking a region encompassing the *tde1-tdi1, vgrG1* genes, and the *vgrG2* operon harboring *tde2-tdi2*. As expected Hcp did not accumulate in the extracellular fraction of this mutant. However, when overexpressing wild type VgrG1 or even VgrG1 truncated variants (812, 784, and 785 a.a.) that are abrogated in their ability to interact with Tde1, Hcp could be detected at high levels in the extracellular fraction (Fig. S3). The shortest variant of VgrG1 (781 a.a.) is previously shown to be incapable of restoring Hcp secretion in Δ*G1*Δ*G2*^16^ and was similarly unable to restore Hcp secretion in Δ*tde1-tdi1*Δ*G1*Δ*G2op* (Fig. S3). These results suggest that the coordinated expression of T6SS components is necessary and influence the regulation of the T6SS. In conclusions, loading cargo effectors onto cognate VgrG proteins is important for T6SS assembly for ejecting the Hcp tube and VgrG spike proteins, and when both Tde1 and Tde2 are not loaded onto VgrG1 and VgrG2 respectively, T6SS is not efficiently assembled for firing.

We recently demonstrated that several strains of *A. tumefaciens* encode functional T6SSs that are necessary for interbacterial competition (Wu et al., in press). From these, *A. tumefaciens* strains 1D1108, 15955 and 12D1, which belong to different genomospecies with only ~80% gene content similarity to C58, were selected to test whether effector loading onto VgrG is a generalizable mechanism for regulating T6SS in genetically diverse *A. tumefaciens*. These three strains each carry a single module that circumscribe *vgrG* and several downstream genes, named as vgrG-associated genes (*V1-5* for 12D1, *V1-9* for 1D1108, *V1-7* for 15955), which encode putative toxin-immunity pairs (*V3-4* in 12D1, *V6-7* in 1D1108, and *V6-7* in 15955) (Fig. 2a, Wu et al., in press). Mutants lacking genes encoding predicted toxin and/or adaptor/chaperone had substantially reduced levels of Hcp and VgrG proteins, as compared to that of their respective wild type strains (Fig. 2b). The polymutant of strain 15955 included the PAAR-encoding gene. PAAR is a core component of T6SS of *Serratia marcescens, Vibrio cholerae* and *Acinetobacter baylyi*^12,18^. In *A. tumefaciens* strain C58, PAAR appears to have only a minor role ^16^. Nevertheless, we tested whether the deficiency of Hcp secretion in 15955Δ*V1-7* is due to the lack of PAAR. A mutant strain harboring a plasmid with the *paar* gene failed to secrete Hcp to wild type levels, suggesting that the deficiency of Hcp secretion in the polymutant was due to the absence of other genes (Fig. 2b and 2d). Consistent with the secretion results, interbacterial competition showed these mutants were as compromised as their corresponding Δ*tssL* mutants in antagonizing the growth of *E. coli* (Fig. 2c). Data indicate that effector loading is a conserved mechanism for regulating T6SS of *A. tumefaciens* and perhaps also in other bacterial species.

**Fig. 2.**
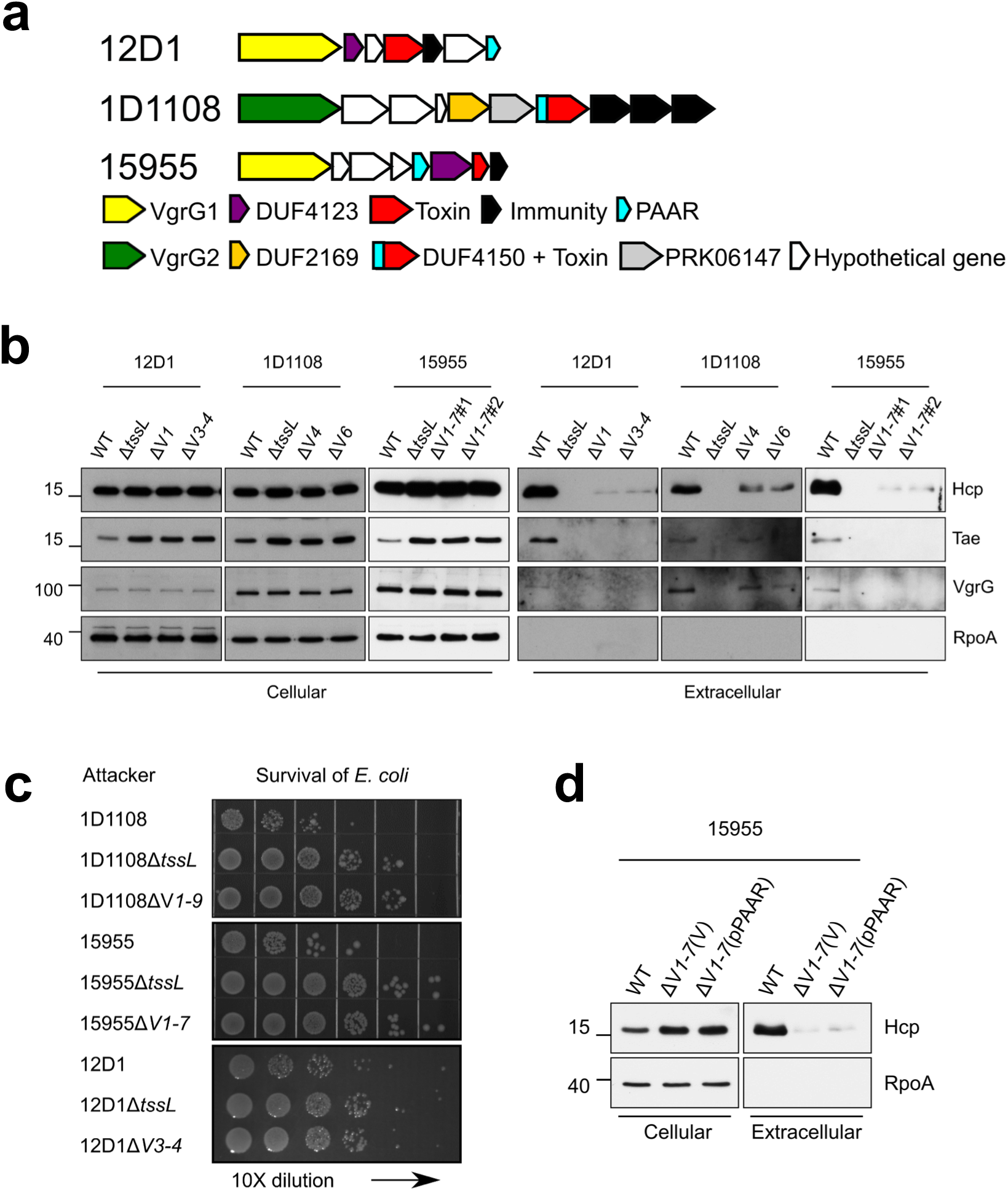
Deletion of vgrG-associated genes leads to decreases in Hcp and VgrG secretion in other *A. tumefaciens* strains. (a) The *vgrG* genetic modules of the tested *A. tumefaciens* strains. Genes are color coded according to the predicted function or results of functional assays (Wu et al., in press). The vgrG-associated genes are *V1-V6* for 12D1, *V1-V9* for 1D1108, *V1-V7* for 15955. Wild type and various mutants of 12D1, 1D1108 and 15955 were analyzed for secretion (b) and antibacterial activity (c) in a ratio of 30:1 against *E. coli* harboring the plasmid pRL662. The target *E. coli* cells were serially-diluted and grown overnight on gentamicin-containing LB agar prior to photographing. Each competition was done at least four times and in three independent experiments. (d) Trans complementation of *paar* cannot restore Hcp secretion in 15955 Δ*V1-7* mutant. T6SS secretion assay of *A. tumefaciens* 15955 Δ*V1-7* containing a pRL662 vector (V) or its derivative expressing *paar* (pPAAR). Cellular and extracellular fractions were collected from *A. tumefaciens* strains grown in liquid 523 medium. Proteins were collected and analyzed in western blots probed with indicated antibodies. RpoA is RNA polymerase and located in the cytosol of *A. tumefaciens*. Molecular weight markers (in kDa) are indicated on the left.

Given these findings, we next tested whether the presence of Tde effectors is critical to initiate Hcp polymerization and assembly of the TssBC sheath. TssB fused with a fluorescent protein has been shown to assemble into the TssBC sheaths of different lengths that were observed as fluorescent foci (short sheaths) or streaks (extended sheaths) across the cell width ^6,19^ In addition, the sheaths were reported to be dynamical contractile structures that were able to contract and disassemble after the extended sheaths were formed ^6,19^ Thus, these fluorescent structures under the microscope can be used as an indicator of the sheath formation. Therefore, a plasmid expressing TssB with a C-terminal fusion to green fluorescence protein (TssB-GFP) was expressed in the Δ*tssB*, Δ*tssL*Δ*tssB* and Δ*tdei*Δ*tssB* mutant strains of *A. tumefaciens* C58. TssB-GFP in Δ*tssB* partially restored T6SS-mediated secretion, as determined on the basis of the amount of secreted Hcp relative to that of the Δ*tssB* expressing an unmodified variant of TssB (Fig. S4). The protein abundance of TssB-GFP in different strains were similar by comparing the band intensity of TssB-GFP and its truncated form on western blots (Fig. S4). Similar to the previous reports ^6^, the sheaths of different length were often observed as foci or streaks across the cell width. Moreover, these fluorescent structures were observed in the vast majority of the Δ*tssB* (TssB-GFP) cells (Fig. 3a). We then counted the number of fluorescent structures in each strain (Fig. 3b). The results showed that the ΔtssB(TssB-GFP) strain had slightly more than one fluorescent structure per cell, and the ΔtssBΔtdei(TssB-GFP) strain had very few fluorescent structures (2~5 foci out of 100 cells). The fluorescent streaks were rarely seen in the later strain (Fig. 3a and 3b). In contrast, no streaks were found in the negative control strain, ΔtssBΔtssL(TssB-GFP), if any GFP foci were observed (Fig. 3a and 3b).

**Fig. 3.**
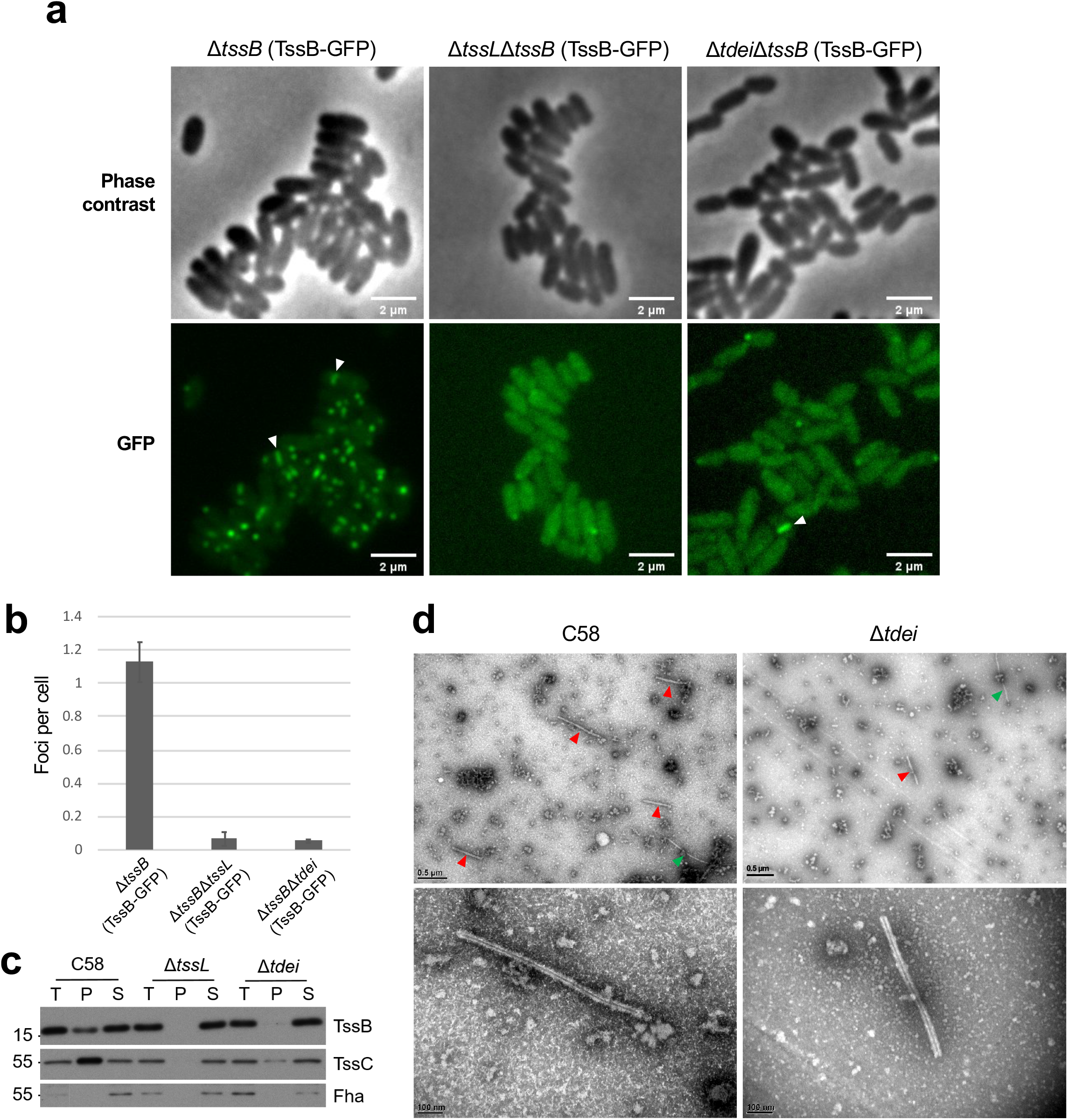
Effector loading onto VgrG is required for efficient TssBC sheath assembly. (a) Observation of TssB-GFP in *A. tumefaciens* C58 strains Δ*tssB, ΔtssLΔtssB*, and Δ*tdei*Δ*tssB* cells each expressing TssB-GFP from pTrc200. Upper panel: phase contrast images; Lower panel: fluorescence images. GFP streaks were indicated by arrows. (b) Quantification of TssB-GFP foci in different genetic background. The number of the fluorescent foci were counted using the “analysis particle” function in Fiji with a manually set threshold. The number of cells were counted manually. For each strain, a total of three images were obtained from independent experiments and each image contained more than 300 cells for quantification and statistical analysis. (c) Western blots of the isolated sheath, which were prepared via ultracentrifugation of samples from *A. tumefaciens* C58 wild type, Δ*tssL*, and Δ*tdei*. T: total proteins from the cell lysate. P: pellet samples containing TssBC sheaths after ultracentrifugation. S: supernatant after ultracentrifugation. Proteins were analyzed in western blots probed with indicated antibodies. Fha, a cytoplasmic protein, serves as a control. Molecular weight markers (in kDa) are indicated on the left. Similar results were obtained from at least two independent experiments. (d) Pellet samples of wild type and Δ*tdei* were visualized with transmission electron microscopy (TEM). Sheaths are indicated by red arrow heads and flagella, which are distinguishable on the basis of smoother, more solid, and thinner structures (~13 nm in width), are indicated by green arrow heads ^27^.

We also isolated and characterized T6SS sheaths, which are tubular structures formed by TssB and TssC proteins ^6^. Pellet and soluble fractions from crude protein extracts of the wild type, Δ*tssL* and Δ*tdei* strains of *A. tumefaciens* C58 were first analyzed by western blots (Fig. 3c). The presence of TssB and TssC proteins in the pellet indicates that they have polymerized into a sheath, whereas monomers/non-polymerized subunits remain in the supernatant. TssB and TssC proteins were detected in the pellet fraction of the wild type but not of the Δ*tssL* mutant. The amounts of TssB and TssC detected in the pellet fractions of Δ*tdei* were substantially lower than those detected in the wild type, suggesting that the TssBC sheaths were inefficiently formed in the absence of Tde effector-encoding genes.

The pellet fractions were further characterized via transmission electron microscopy (TEM). Sheath-like structures were easily detected in fractions from C58 and only very few were observed in fractions from Δ*tdei* (Fig. 3d). No sheath structures were identified from the fraction from Δ*tssL* (not shown). Regardless of the source, the sheaths were similar in structure. The diameter of the sheath structure was calculated to be ~30 nm. They have a hollow lumen, suggesting these were contracted sheathes which had ejected the Hcp tube. Relative to sheaths of other bacteria, those from *A. tumefaciens* have a similar morphology and diameter with the ones reported in other bacteria (25-33 nm) ^6,20–23^. The data together strongly suggest that effector-loaded VgrG is an important trigger for efficient TssBC sheath assembly. Such a mechanism may be employed by *A. tumefaciens* to prevent the assembly of the T6SS machine when effectors are not present and/or loaded.

One important question raised from this study is whether regulation of the T6SS via effector loading onto VgrG is a widespread mechanism. In many other species of bacteria, such a mechanism may have been overlooked because of the lack of mutants deleted of all effector-encoding genes and the difficulties to identify VgrG variants that uncouple the two functions of VgrG, those being the delivery of effectors and being a structural feature of the T6SS. Our findings contributed to a new model that predicts the loading of effector cargo onto their cognate VgrG spike protein regulates assembly and activation of the T6SS.

Previous studies provided evidence that VgrG interacts with components of the baseplate and the interactions are critical for assembly of the T6SS ^7,8,24,25^ Our findings further suggest that effector loading onto VgrG spike is the key in completing T6SS assembly. It is possible that effector-loaded VgrG exhibits higher affinity than VgrG itself for recruitment onto the membrane-associated baseplate. Alternatively, effector loading onto the VgrG-baseplate complexes is the trigger for Hcp polymerization and/or TssBC sheath assembly. In conclusion, our study reveals a mechanism that ensures effectors are loaded onto T6SS prior to completing its assembly and firing outwards. Such a mechanism may be deployed by not only *A. tumefaciens* but also other T6SS-possessing bacteria to regulate a system that influences their fitness and composition of their communities.

## Materials and Methods

### Bacterial strains, growth conditions, and molecular techniques

Bacterial strains and sequences of primers used in this study are listed in Tables S1 and S2, respectively. *A. tumefaciens* was grown in 523 medium at 25 °C and *E. coli* was grown in LB medium at 37 °C, unless otherwise indicated ^10,16,17^ Antibiotics and concentrations used were: gentamycin (50 μg/mL for *A. tumefaciens* and 30 μg/mL for *E. coli)* and spectinomycin (200 μg/mL). Procedures for preparing DNA, PCR, and cloning are described in SI.

### Secretion assay

Secretion assays were performed as described ^16^. Briefly, *A. tumefaciens* strains were cultured in 523 medium overnight and sub-cultured, using an initial cell density of OD_600nm_ = 0.2, in I-medium (pH 5.5) or 523 medium, depending on the design of the experiments. After 6 hours of subculturing, the samples were centrifuged at 10,000 g for 10 min to separate the extracellular and the cellular fractions. The cell pellets were adjusted to OD_600nm_ = 5.0 and the respective extracellular fractions were filtered through low protein-binding 0.22 μm sterilized filter units (Milipore, Tullagreen, Ireland) and proteins were precipitated in trichloroacetic acid (TCA) ^15^. Western analyses were done as previously described ^10,16,17^

### Interbacterial competition assay

Methods for interbacterial competition assays were previously described ^10^. Briefly, *A. tumefaciens* strains were co-cultured at with *E. coli* K-12 cells harboring the plasmid pRL662 (confers gentamycin resistance), at a ratio of 30:1 on LB agar. The surviving *E. coli* cells were serially diluted, spotted or quantified by counting colony forming units (CFUs) on gentamycin-containing LB agar plates. Statistics were calculated using one-way ANOVA and Tukey’s honestly significance difference (HSD) test (http://astatsa.com/OneWay_Anova_with_TukeyHSD/).

### Sheath preparation and transmission electron microscopy (TEM)

Isolation of sheath preparations was performed by following methods previously described ^6^. In brief, *A. tumefaciens* cells were cultured overnight in 5 mL 523 and subcultured into 50 mL I-medium at 25 °C for 6 hours. The cells were harvested and lysed for 15 mins at 37 °C in 4 mL of buffer containing 0.5X CelLytic B (Sigma-Aldrich, St. Louis, USA), 150 mM NaCl, 50 mM Tris pH 7.4, lysozyme (500 μg/mL), DNase I (50 μg/mL), 1 mM phenlmethylsulphonyo-fluride (PMSF) and 0.05%Triton X-100. Cell debris was removed via centrifugation at 10,000 g, 10 mins at 4 °C. The high-molecular-weight complexes were separated from the clear cell lysate via ultracentrifugation at 150,000 g for 1 hour at 4 °C. Pellet fractions were resuspended in 150 μL buffer containing 150 mM NaCl, 50 mM Tris pH7.4 and 0.75 μL Protease Inhibitor Cocktail Set III (EMD Millipore, Pacific Center Court San Diego, USA). The samples were analyzed via western analyses. For TEM, 10 μL of the samples were deposited on copper grids with carbon-formvar film support. After 3 minutes, samples were negative stained for 90 seconds with 25% uranyl acetate. A Tecnai G2 Spirit TEM (FEI, Hillsboro, USA) set at 80 kV was used to visualize samples.

### Fluorescence microscopy

Cells were cultured in 523 medium overnight followed by subculturing in I-medium for 3 hours. One mL of each bacterial culture was fixed with 8 μL 25 % glutaraldehyde and 125 μL 37 % formaldehyde for 20 minutes and washed twice with PBS (Biomate, Taiwan) with additional 0.5 % Tween 20 (PBST). Bacterial cells were resuspended with 30 μL PBST. 3 μL of the fixed cell suspension was deposited onto agarose pad (PBST with 2% agarose) to reduce random movement of cells. An upright microscope BX61 (Olympus, Tokyo, Japan) equipped with a CMOS camera (C11440 ORCA-R2 Flash 2.8, Hamamatsu, Japan), objective lens (UPlanFLN 100x/1.30, Olympus, Tokyo, Japan) and a GFP filter set (Part number: FITC-3540C-000, Semwork, New York, USA) were used. Images were acquired and processed using Improvision Volocity 6.3 software (Perkin Elmer, Waltham, USA) and Fiji ^26^, respectively.

## Acknowledgements

The authors thank Martin Pilhofer and Romain Kooger in the Institute of Molecular Biology & Biophysics, ETH Zürich for their discussions and providing the protocol for preparing TssBC sheaths. We acknowledge the assistance by Claudia Parada of the Institute of Biological Chemistry, and staff in the Plant Cell Biology Core Laboratory as well as DNA Sequencing Core Laboratory at the Institute of Plant and Microbial Biology, Academia Sinica, Taiwan. The authors also thank Lay-Sun Ma, Chih-Horng Kuo, and Romain Kooger for critically reading the manuscript and the members of Lai laboratory for their discussions and suggestions. Funding for this project was provided by the Ministry of Science and Technology of Taiwan (MOST) (grant no. 104-2311-B-001-025-MY3 to E.-M.L and 106-2311-B-001-009 to Y.-L.S). Work in the Chang lab is supported in part by the National Institute of Food and Agriculture, US Department of Agriculture award 2014-51181-22384. The funders had no role in study design, data collection and interpretation, or the decision to submit the work for publication.

## Author Contributions

YWL, CFW, and EML designed and conceived the experiments. YWL, CFW, and JSL performed the experiments. DB and YLS provided the tools. JHC, YLS, and EML supervised the execution of the experiments. YWL, EML, and JHC, with contributions from CFW, JSL, and YLS, wrote the paper. All authors read and approved the paper.

## SI EXPERIMENTAL PROCEDURE

### DNA preparation and plasmids construction

Plasmid DNA was extracted using Presto Mini Plasmids Kit (Geneaid, Taiwan). 2X Manufacturer instructions were followed in using Ready Mix A (Zymeset, Taiwan) for polymerase chain reactions (PCRs). For construction of pTrc-TssB-GFP, *A. tumefaciens tssB* and the upstream sequences corresponding to the ribosome binding site were amplified from pTrc-TssB (EML4043) (Table S1) with primers tssB_XmaI_F and *tssB-GFP*_R (Table S2). The *gfp* gene was amplified from pBBR1-GFP (EML3) ^1^ using primers GFP_HindIII_R and *tssB-GFP*_F. The two amplified products were fused together in a second PCR that used primers tssB_XmaI_F and GFP_HindIII_R, which include restriction site sequences for XmaI and HindIII. The purified fusion product and pTrc200 plasmid were digested with XmaI and HindIII-HF (New England BioLabs, Ipswich, USA) and subsequently ligated together using T4 DNA ligase (New England BioLabs, Ipswich, USA). The plasmid construct was confirmed via colony PCR, enzyme digestion, sequencing and western blot of *A. tumefaciens* cells harboring the plasmid.

### Mutant construction

The pJQ200KS suicide plasmid ^2^ and double crossover method were used to generate in-frame deletions of *A. tumefaciens* genes ^3^. In brief, cells were electroporated with suicide plasmids, transformants were selected on 523 agar plates containing gentamycin without sucrose. The Gm-resistant colonies were cultured overnight in LB broth without Gm, serially diluted and spread onto 523 agar plates containing 5% sucrose without Gm to enrich for bacterial cells that had undergone a second crossover event. The deletion mutants were confirmed via colony PCR and western blot analyses.

**Fig. S1.**
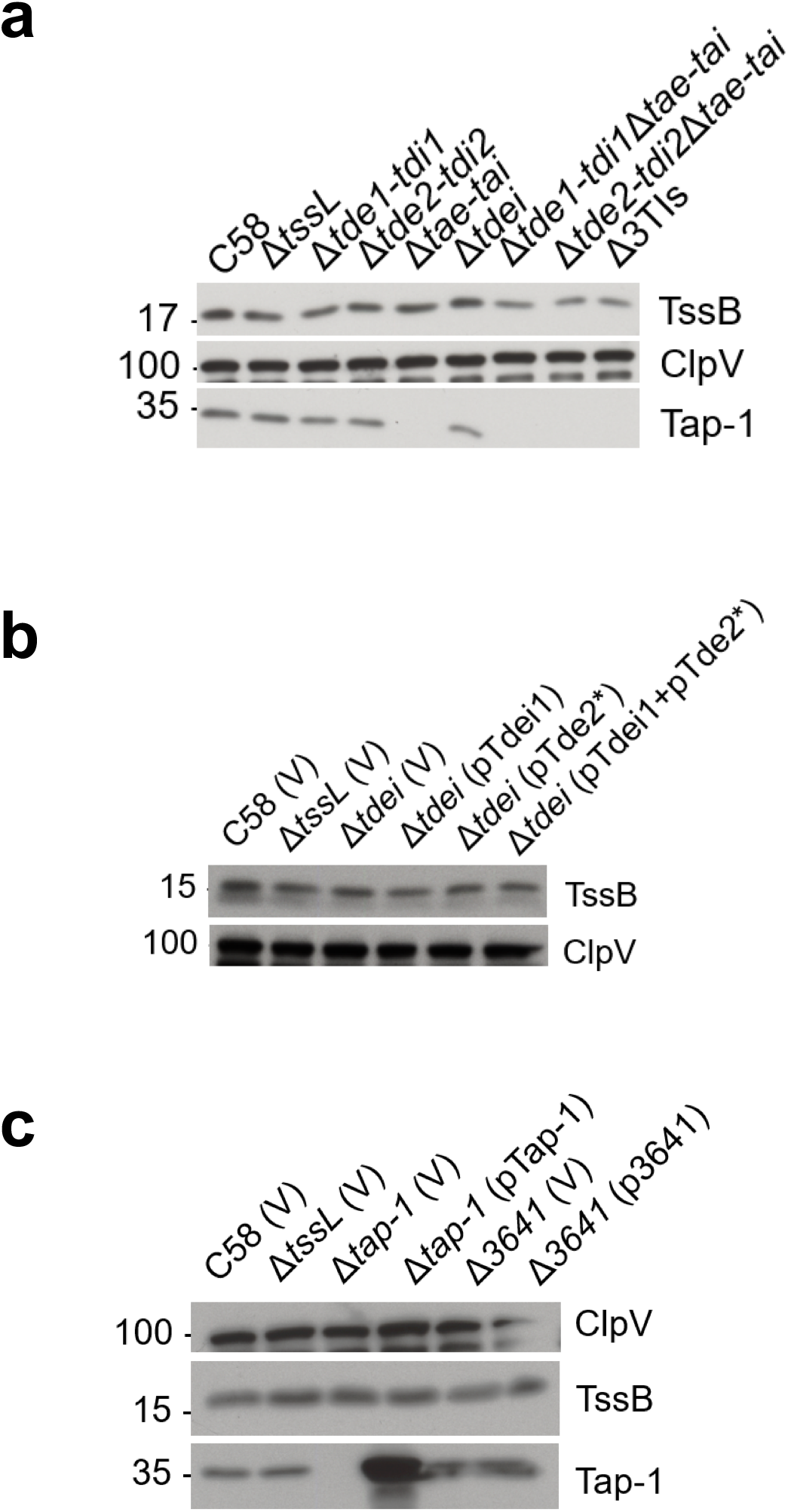
Western blots of cellular fractions of representative T6SS proteins from Figure 1. Cellular fractions were collected from *A. tumefaciens* strains grown in liquid 523 medium. (a) same cellular fractions of Fig. 1b; (b) same cellular fractions of Fig. 1c; (c) same cellular fractions of Fig. 1e. Western blots were probed with antibodies against TssB (representative protein encoded by *imp* operon), ClpV (representative protein encoded by *hcp* operon), and Tap-1. Molecular weight markers (in kDa) are indicated on the left.

**Fig. S2.**
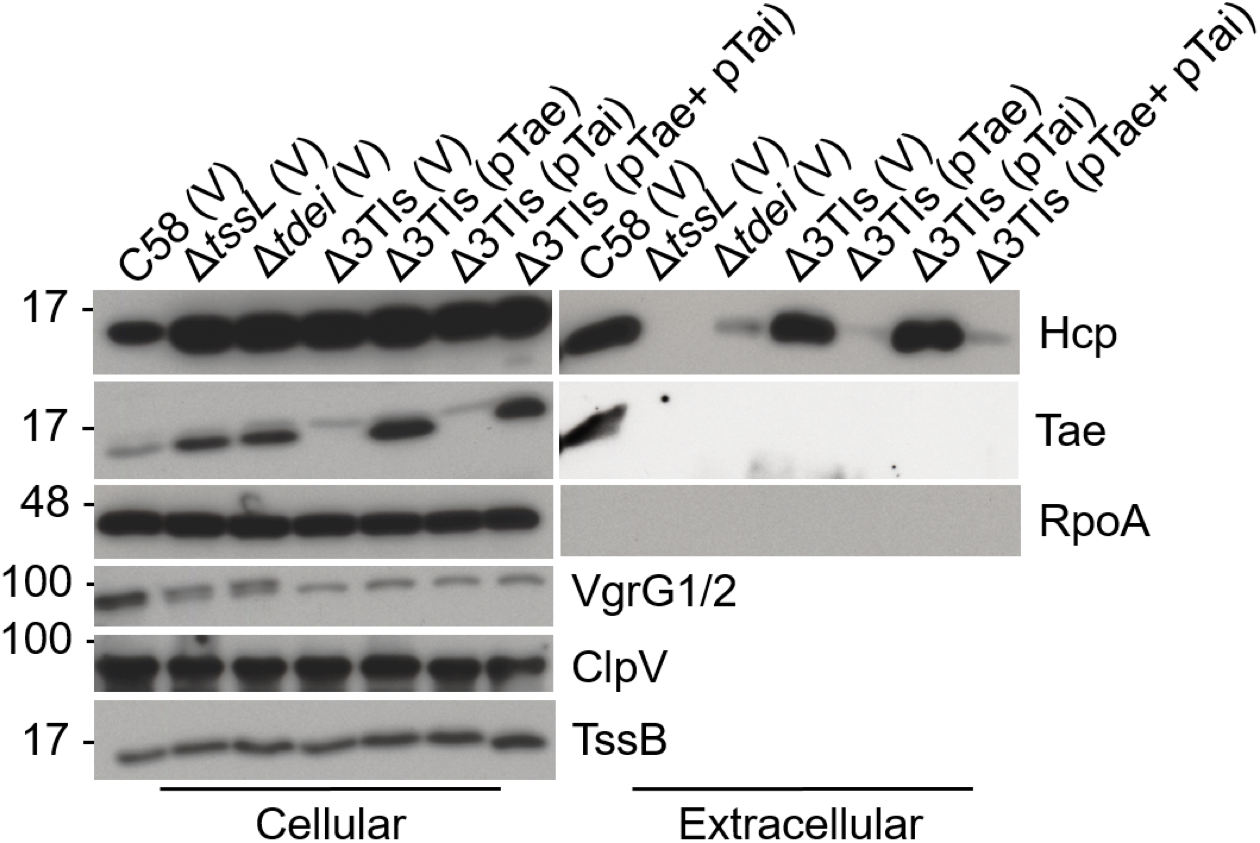
Effect of *tae* on Hcp secretion in the Δ*tdei* mutant. *A. tumefaciens* wild type C58, Δ*tssL*, Δ*tdei*, and Δ3TIs harboring respective plasmids were used. *A. tumefaciens* cells were grown in I-medium (pH 5.5) and cellular and extracellular fractions were collected for western blot analysis using various antibodies as indicated. Plasmids: pRL662 vector (V), pTae (*tae* expressed pRL662), pTai (*tai* expressed on pTrc200). Molecular weight markers (in kDa) are indicated on the left.

**Fig. S3.**
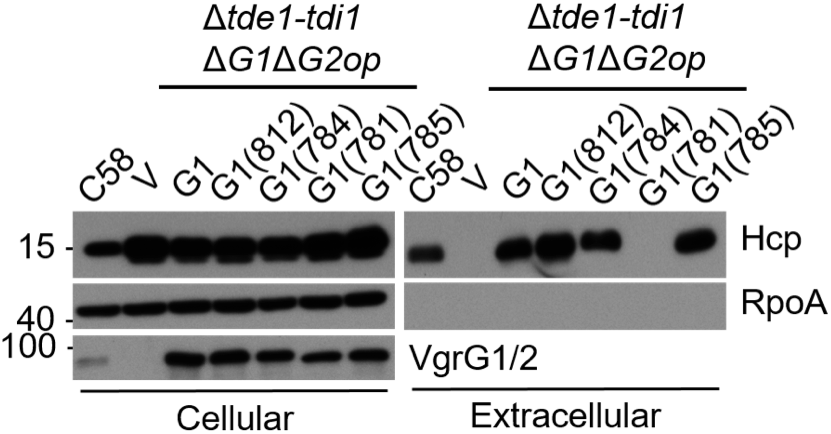
Overexpression of functional VgrG1 variants is able to restore Hcp secretion in the absence of Tde effectors. *A. tumefaciens* C58 Δ*tdei*Δ*vgrG1* Δ*vgrG2* (Δ*tdei*Δ*G1*Δ*G2op* mutant) harboring a pRL662 vector (V) or its derivatives overexpressing full-length and truncated VgrG1 proteins were used. *A. tumefaciens* cells were grown in I-medium (pH 5.5) and cellular and extracellular fractions were collected for western blot analysis using various antibodies as indicated. Molecular weight markers (in kDa) were indicated on the left.

**Fig. S4.**
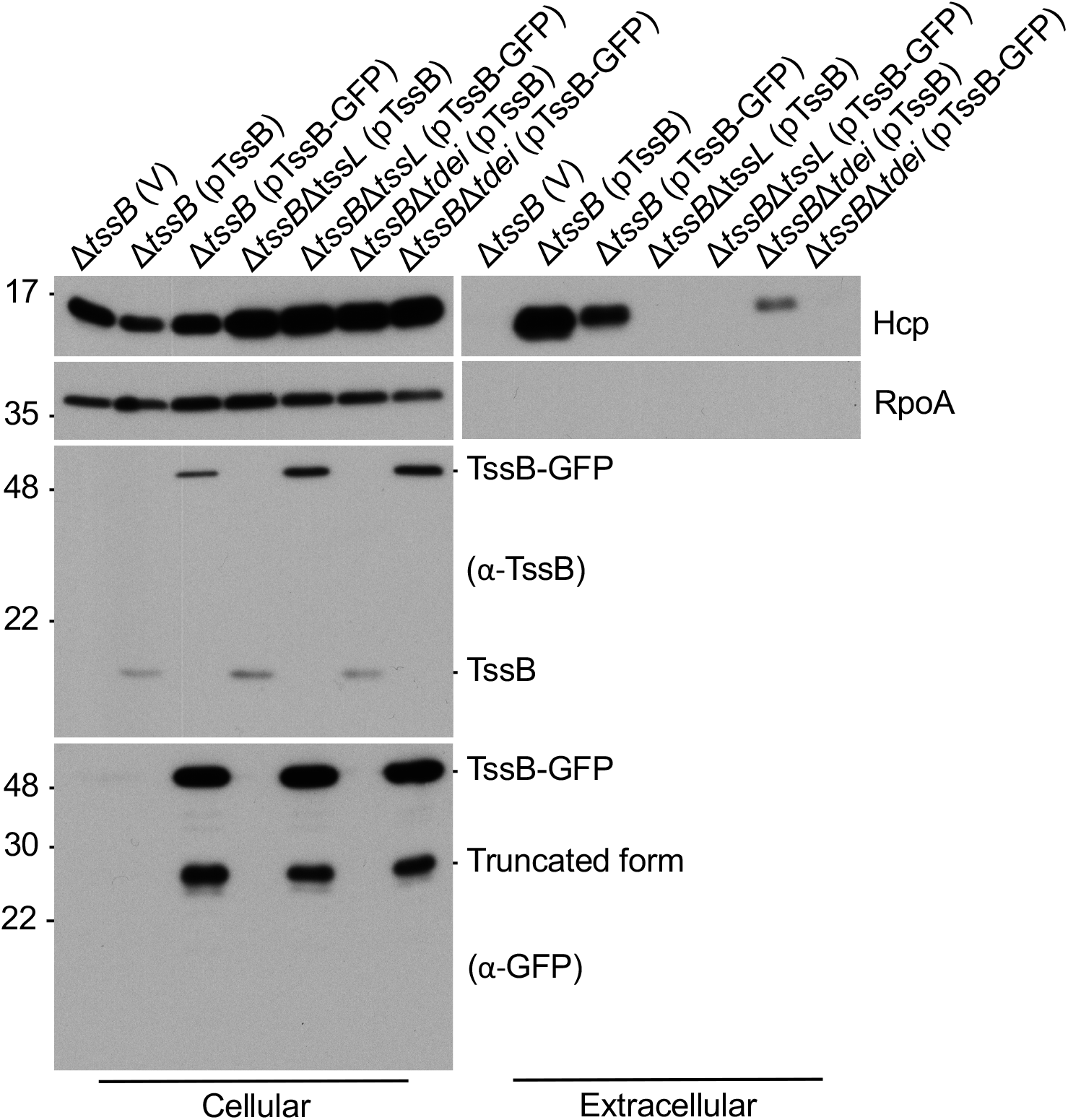
C-terminal GFP-tagged TssB partially complemented Hcp secretion in Δ*tssB*. Various *A. tumefaciens* strains were grown in I-medium (pH5.5) and the cellular and extracellular (S) fractions were collected for western blotting using various antibodies as indicated. Molecular weight markers (in kDa) were indicated on the left.

**Table S1.**
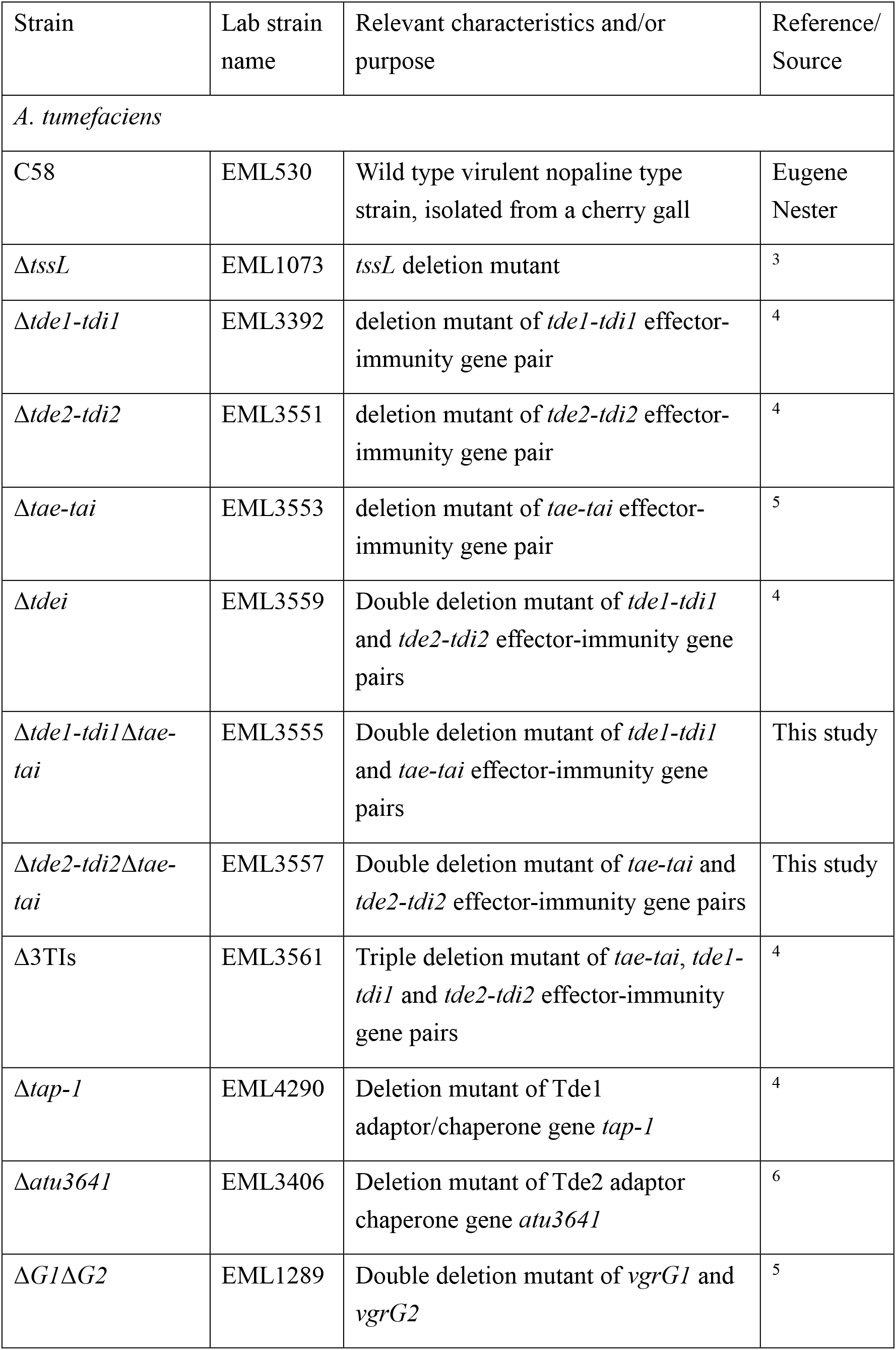
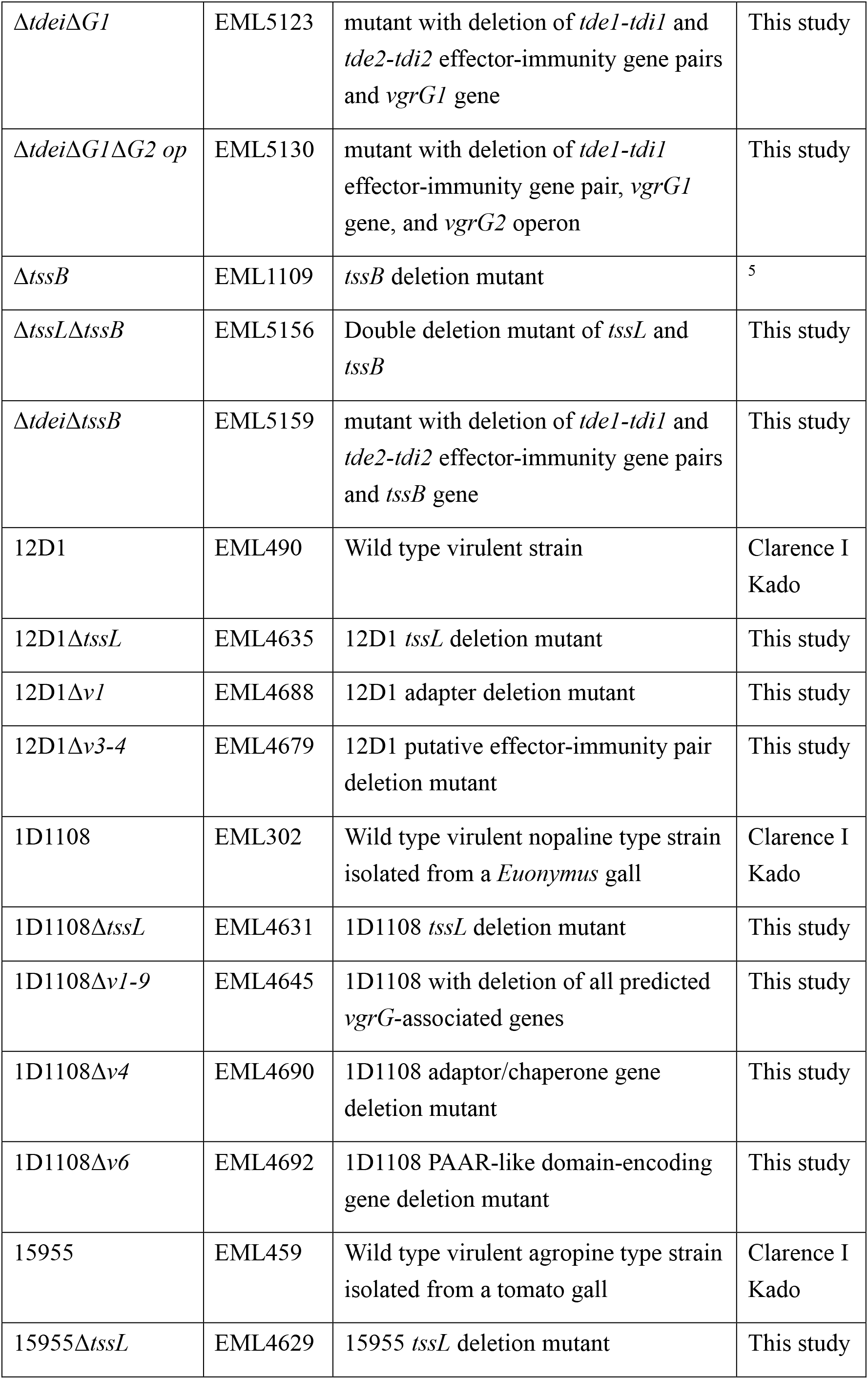
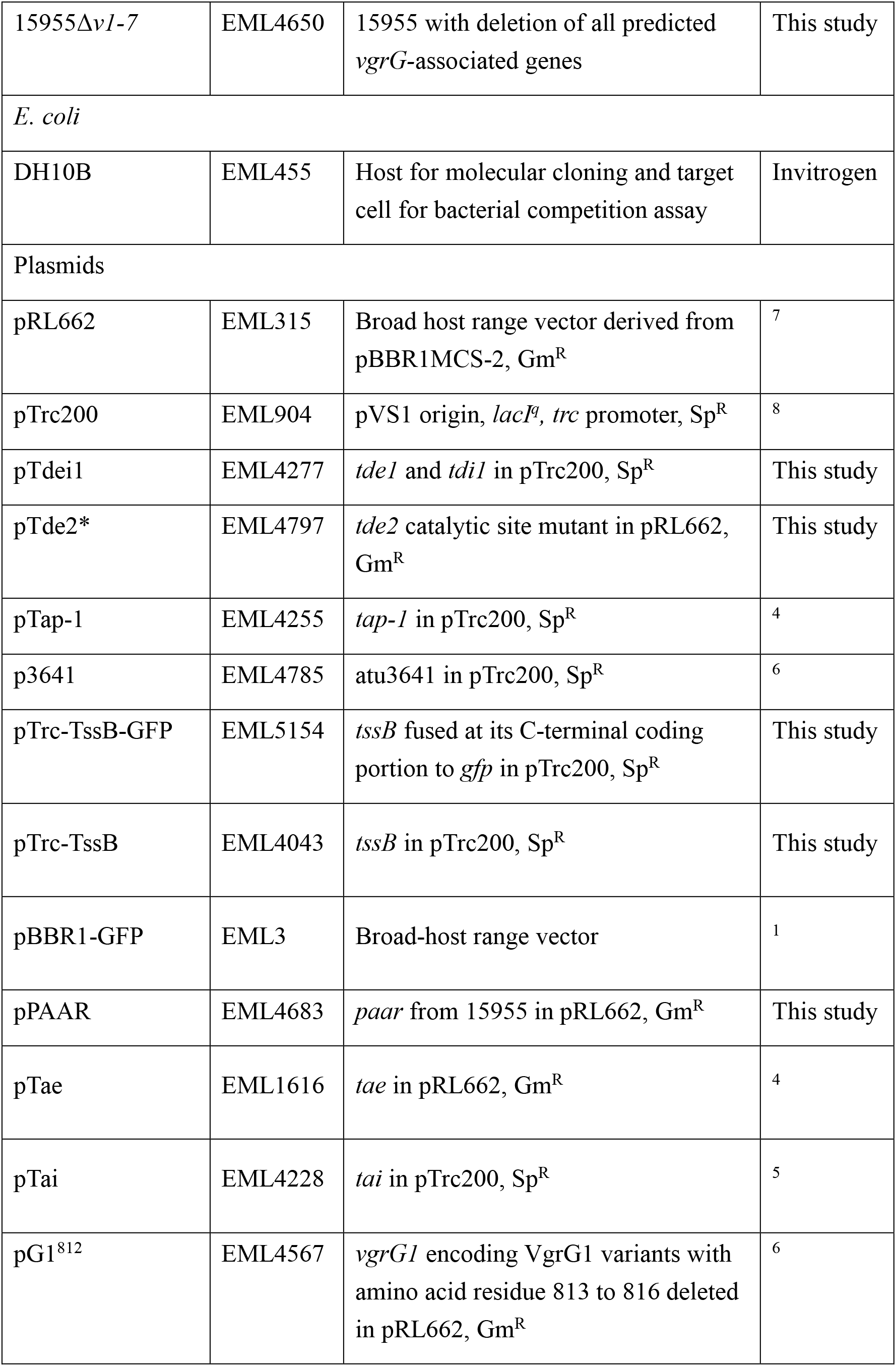
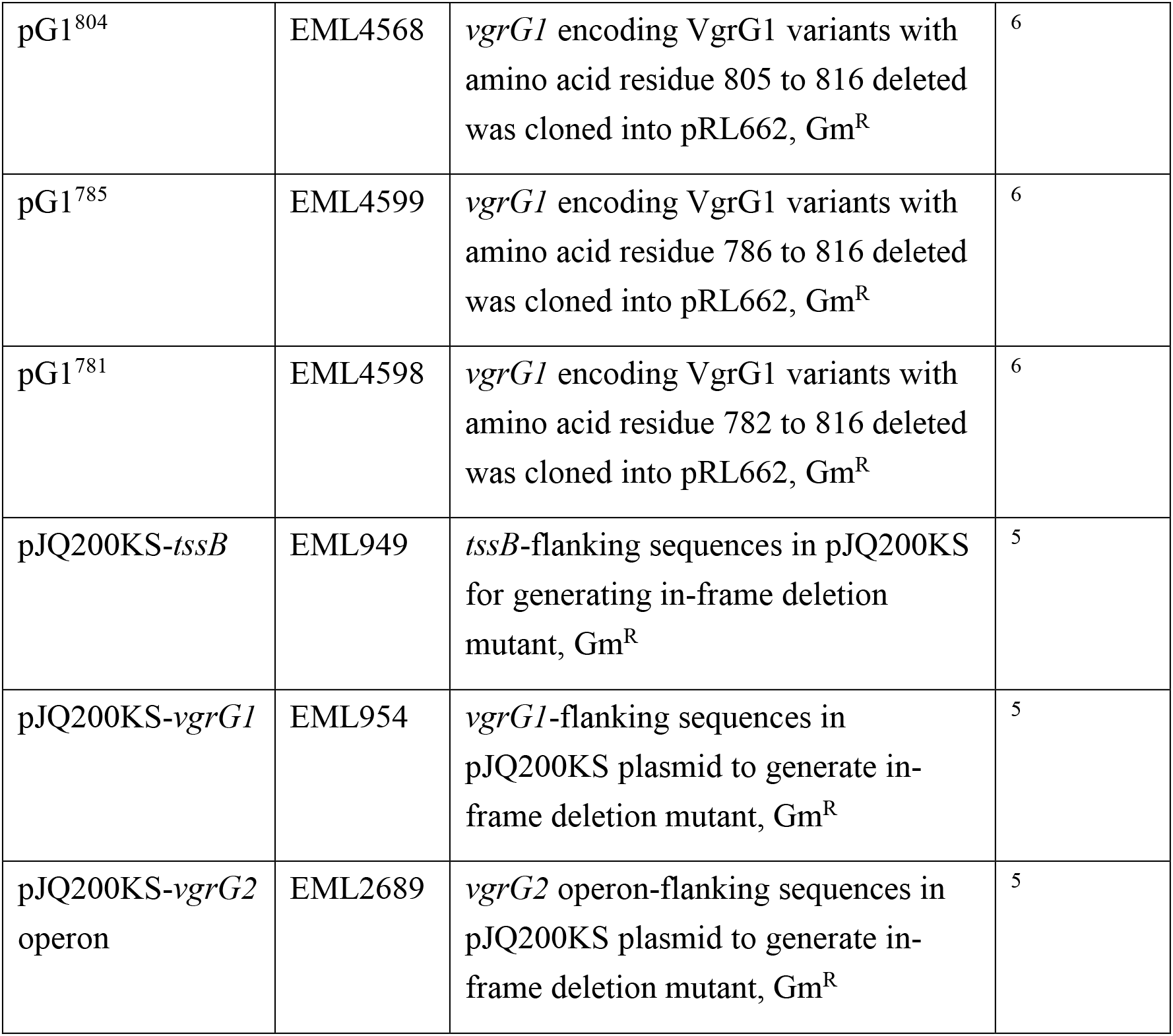
Strains and plasmids used in this study.

**Table S2.**
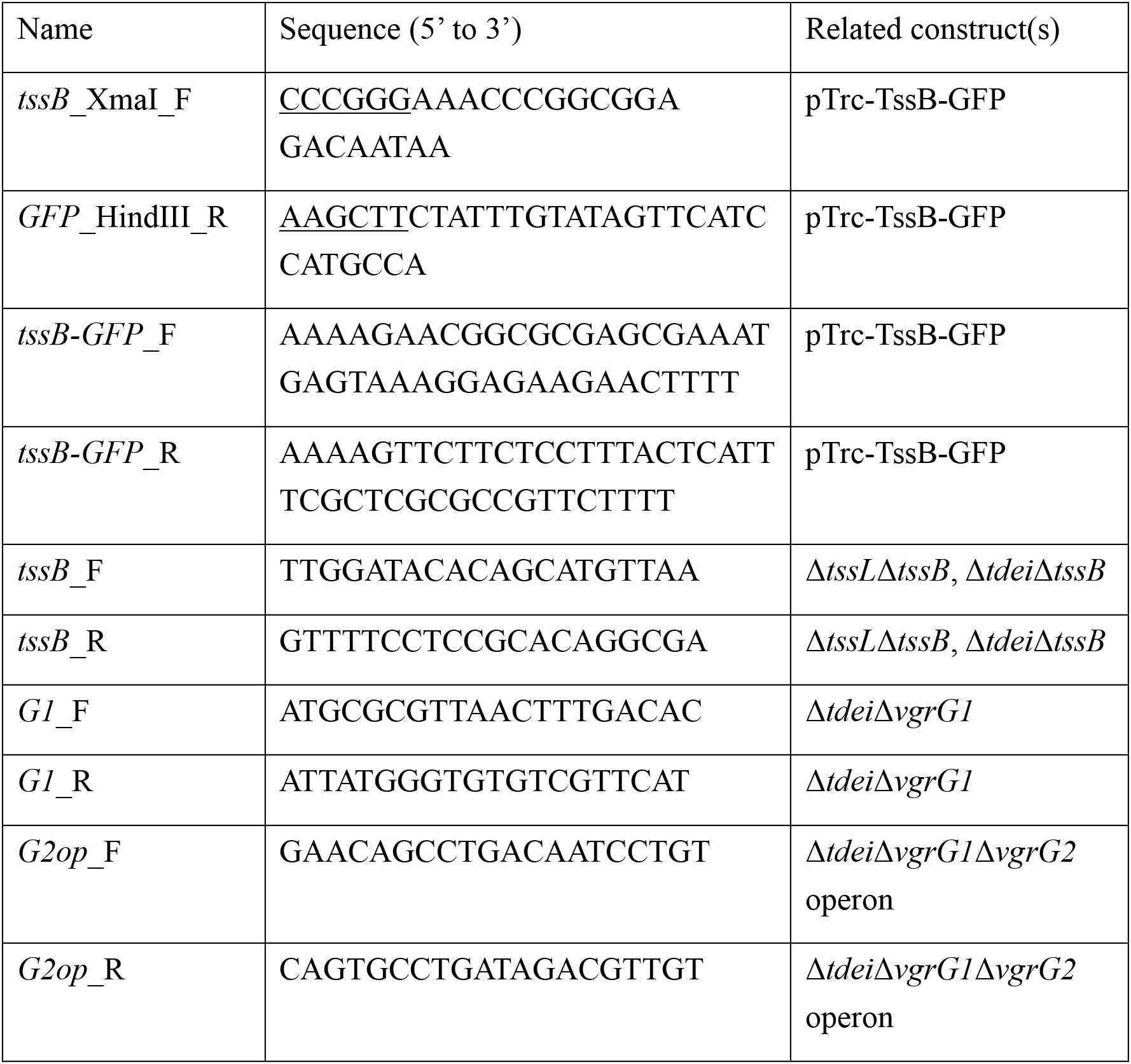
Primers used in this study.

## REFERENCES

1 Russell, A. B., Peterson, S. B. & Mougous, J. D. Type VI secretion effectors: poisons with a purpose. Nature reviews. Microbiology 12, 137–148, doi:10.1038/nrmicro3185 (2014).

2 Lien, Y.-W. & Lai, E.-M. Type VI Secretion Effectors: Methodologies and Biology. Frontiers in Cellular and Infection Microbiology 7, 254, doi:10.3389/fcimb.2017.00254 (2017).

3 Ringel, P. D., Hu, D. & Basler, M. The Role of Type VI Secretion System Effectors in Target Cell Lysis and Subsequent Horizontal Gene Transfer. Cell reports 21, 3927–3940, doi:10.1016/j.celrep.2017.12.020 (2017).

4 Bernal, P., Llamas, M. A. & Filloux, A. Type VI secretion systems in plant-associated bacteria. Environmental microbiology 20, 1–15, doi:10.1111/1462-2920.13956 (2018).

5 Verster, A. J. et al. The Landscape of Type VI Secretion across Human Gut Microbiomes Reveals Its Role in Community Composition. Cell host & microbe 22, 411–419.e414, doi:10.1016/j.chom.2017.08.010 (2017).

6 Basler, M., Pilhofer, M., Henderson, G. P., Jensen, G. J. & Mekalanos, J. J. Type VI secretion requires a dynamic contractile phage tail-like structure. Nature 483 (2012).

7 Brunet, Y. R., Zoued, A., Boyer, F., Douzi, B. & Cascales, E. The Type VI Secretion TssEFGK-VgrG Phage-Like Baseplate Is Recruited to the TssJLM Membrane Complex via Multiple Contacts and Serves As Assembly Platform for Tail Tube/Sheath Polymerization. PLoS Genetics 11, e1005545, doi:10.1371/journal.pgen.1005545 (2015).

8 Zoued, A. et al. Priming and polymerization of a bacterial contractile tail structure. Nature 531, 59–63, doi:10.1038/nature17182 (2016).

9 Cianfanelli, F. R., Monlezun, L. & Coulthurst, S. J. Aim, Load, Fire: The Type VI Secretion System, a Bacterial Nanoweapon. Trends in microbiology 24, 51–62, doi:10.1016/j.tim.2015.10.005 (2016).

10 Ma, L.-S., Hachani, A., Lin, J.-S., Filloux, A. & Lai, E.-M. Agrobacterium tumefaciens Deploys a Superfamily of Type VI Secretion DNase Effectors as Weapons for Interbacterial Competition In Planta. Cell host & microbe 16, 94–104, doi:10.1016/j.chom.2014.06.002 (2014).

11 Flaugnatti, N. et al. A phospholipase A1 antibacterial Type VI secretion effector interacts directly with the C-terminal domain of the VgrG spike protein for delivery. Molecular Microbiology 99, 1099–1118, doi:10.1111/mmi.13292 (2016).

12 Cianfanelli, F. R. et al. VgrG and PAAR Proteins Define Distinct Versions of a Functional Type VI Secretion System. PLoS Pathogens 12, e1005735, doi:10.1371/journal.ppat.1005735 (2016).

13 English, G. et al. New secreted toxins and immunity proteins encoded within the Type VI secretion system gene cluster of Serratia marcescens. Molecular Microbiology 86, 921–936, doi:10.1111/mmi.12028 (2012).

14 Hood, R. D. et al. A Type VI Secretion System of Pseudomonas aeruginosa Targets a Toxin to Bacteria. Cell host & microbe 7, 25–37, doi:10.1016/j.chom.2009.12.007 (2010).

15 Wu, H.-Y., Chung, P.-C., Shih, H.-W., Wen, S.-R. & Lai, E.-M. Secretome Analysis Uncovers an Hcp-Family Protein Secreted via a Type VI Secretion System in Agrobacterium tumefaciens. Journal of Bacteriology 190, 2841–2850, doi:10.1128/JB.01775-07 (2008).

16 Bondage, D. D., Lin, J.-S., Ma, L.-S., Kuo, C.-H. & Lai, E.-M. VgrG C terminus confers the type VI effector transport specificity and is required for binding with PAAR and adaptor-effector complex. Proceedings of the National Academy of Sciences of the United States of America 113, E3931–E3940, doi:10.1073/pnas.1600428113 (2016).

17 Lin, J.-S., Ma, L.-S. & Lai, E.-M. Systematic Dissection of the Agrobacterium Type VI Secretion System Reveals Machinery and Secreted Components for Subcomplex Formation. PLoS ONE 8, e67647, doi:10.1371/journal.pone.0067647 (2013).

18 Shneider, M. M. et al. PAAR-repeat proteins sharpen and diversify the Type VI secretion system spike. Nature 500, 350–353, doi:10.1038/nature12453 (2013).

19 Basler, M. & Mekalanos, J. J. Type 6 secretion dynamics within and between bacterial cells. Science (New York, N.Y.) 337, 815–815, doi:10.1126/science.1222901 (2012).

20 Lossi, N. S. et al. The HsiB1C1 (TssB-TssC) Complex of the Pseudomonas aeruginosa Type VI Secretion System Forms a Bacteriophage Tail Sheathlike Structure. The Journal of Biological Chemistry 288, 7536–7548, doi:10.1074/jbc.M112.439273 (2013).

21 Wang, J. et al. Cryo-EM structure of the extended type VI secretion system sheath-tube complex. Nature microbiology 2, 1507–1512, doi:10.1038/s41564-017-0020-7 (2017).

22 Salih, O. et al. Atomic Structure of Type VI Contractile Sheath from Pseudomonas aeruginosa. Structure 26, 329–336.e323, doi:10.1016/j.str.2017.12.005 (2018).

23 Clemens, D. L., Ge, P., Lee, B. Y., Horwitz, M. A. & Zhou, Z. H. Atomic structure of T6SS reveals interlaced array essential to function. Cell 160, 940–951, doi:10.1016/j.cell.2015.02.005 (2015).

24 Planamente, S. et al. TssA forms a gp6-like ring attached to the type VI secretion sheath. The EMBO journal 35, 1613–1627, doi:10.15252/embj.201694024 (2016).

25 Dix, S. R. et al. Structural insights into the function of type VI secretion system TssA subunits. Nature Communications 9, 4765, doi:10.1038/s41467-018-07247-1 (2018).

26 Schindelin, J. et al. Fiji: an open-source platform for biological-image analysis. Nature Methods 9, 676, doi:10.1038/nmeth.2019 (2012).

27 Lai, E. M., Chesnokova, O., Banta, L. M. & Kado, C. I. Genetic and environmental factors affecting T-pilin export and T-pilus biogenesis in relation to flagellation of Agrobacterium tumefaciens. Journal of bacteriology 182, 3705–3716 (2000).

## References

1 Ouahrani-Bettache, S., Porte, F., Teyssier, J., Liautard, J.-P. & Köhler, S. pBBR1-GFP: A Broad-Host-Range Vector for Prokaryotic Promoter Studies. BioTechniques 26, 620–622, doi:10.2144/99264bm05 (1999).

2 Quandt, J. & Hynes, M. F. Versatile suicide vectors which allow direct selection for gene replacement in gram-negative bacteria. Gene 127, 15–21 (1993).

3 Ma, L.-S., Lin, J.-S. & Lai, E.-M. An IcmF Family Protein, ImpLM, Is an Integral Inner Membrane Protein Interacting with ImpKL, and Its Walker A Motif Is Required for Type VI Secretion System-Mediated Hcp Secretion in Agrobacterium tumefaciens. Journal of Bacteriology 191, 4316–4329, doi:10.1128/jb.00029-09 (2009).

4 Ma, L.-S., Hachani, A., Lin, J.-S., Filloux, A. & Lai, E.-M. Agrobacterium tumefaciens Deploys a Superfamily of Type VI Secretion DNase Effectors as Weapons for Interbacterial Competition In Planta. Cell host & microbe 16, 94–104, doi:10.1016/j.chom.2014.06.002 (2014).

5 Lin, J.-S., Ma, L.-S. & Lai, E.-M. Systematic Dissection of the Agrobacterium Type VI Secretion System Reveals Machinery and Secreted Components for Subcomplex Formation. PLoS ONE 8, e67647, doi:10.1371/journal.pone.0067647 (2013).

6 Bondage, D. D., Lin, J.-S., Ma, L.-S., Kuo, C.-H. & Lai, E.-M. VgrG C terminus confers the type VI effector transport specificity and is required for binding with PAAR and adaptor–effector complex. Proceedings of the National Academy of Sciences of the United States of America 113, E3931–E3940, doi:10.1073/pnas.1600428113 (2016).

7 Vergunst, A. C. et al. VirB/D4-dependent protein translocation from Agrobacterium into plant cells. Science 290, 979–982 (2000).

8 Schmidt-Eisenlohr, H., Domke, N. & Baron, C. TraC of IncN plasmid pKM101 associates with membranes and extracellular high-molecular-weight structures in Escherichia coli. J Bacteriol 181, 5563–5571 (1999).

